# Karrikin-sensing protein KAI2 is a new player in regulating root growth patterns

**DOI:** 10.1101/195891

**Authors:** Stéphanie M. Swarbreck, Yannick Guerringue, Elsa Matthus, Fiona J. C. Jamieson, Julia M. Davies

**Author notes:** Corresponding author, 01223-748-980.

## Abstract

Roots form highly complex systems varying in growth direction and branching pattern to forage for nutrients efficiently. Here mutations in the KAI2 (KARRIKIN INSENSITIVE) α/β-fold hydrolase and the MAX2 (MORE AXILLARY GROWTH 2) F-box leucine-rich protein, which together perceive karrikins (smoke-derived butenolides), caused alteration in root growth direction (root skewing and waving) of *Arabidopsis thaliana*. This exaggerated root skewing was independent of endogenous strigolactone perception by the D14 α/β-fold hydrolase and MAX2. Thus KAI2/MAX2’s regulation of root growth may be through perception of endogenous KAI2-ligands, which have yet to be identified. Degradation targets of the KAI2/MAX2 complex, SMAX1 (SUPPRESSOR OF MAX2-1) and SMXL6,7,8 (SUPPRESSOR OF MAX2-1-LIKE) are also involved in the regulation of root skewing. Genetic data reveal a new potential target for degradation, as mutation in the SKS3 (SKU5 similar) but not the SKU5/SKS17 root plasma membrane glycoprotein suppresses the exaggerated root skewing induced by the lack of MAX2. In *Arabidopsis thaliana* therefore, the KAI2 karrikin-sensing protein acts to limit root skewing, and we propose a mechanism involving root radial expansion as the mutant’s gravitropic and mechano-sensing responses remained largely unaffected.

## Introduction

Roots grow in complex patterns that are highly relevant to their adaptation to different soil conditions and yet very difficult to investigate in this complex medium. *Arabidopsis thaliana* roots grown vertically on solid medium produce specific surface-dependent growth patterns described as skewing (deviation from vertical) and waving (Roy & Bassham, 2014). Established differences amongst *Arabidopsis* ecotypes suggest that these patterns may reflect an adaptive response relevant to natural soil conditions (Vaughn & Masson, 2011; Schultz *et al.*, 2017).

While root skewing has been widely observed and reported, it is not fully understood and no model akin to that available for the root gravitropic response has been proposed (Darwin & Darwin, 1880; Oliva & Dunand, 2007; Roy & Bassham, 2014). The characterization of *Arabidopsis* mutants has been critical in identifying genetic components that can govern root skewing and waving (Okada & Shimura, 1990; Wang *et al.*, 2011; Shih *et al.*, 2014), which represent the integrated response to gravity, light and contact with the solid medium as the root tip grows on the surface of the agar (Thompson & Holbrook, 2004). A skewing phenotype in mutants impaired in mechano-sensing such as *feronia* (Shih *et al.*, 2014) or *cml24* (Wang *et al.*, 2011) supports a link between root skewing and thigmomorphogenesis (morphological change in response to mechanical stimulation from surface contact).

The role of plant hormones in root skewing and waving is poorly understood but auxins (Okada & Shimura, 1990), ethylene (Buer *et al.*, 2000; 2003), cytokinins (Kushwah *et al.*, 2011) and brassinosteroids (Lanza *et al.*, 2012) are implicated. Little is known of the role of a recently characterized set of phytohormones, strigolactones (SL, Roy & Bassham, 2014) and related smoked-derived butenolides, karrikins (KAR, Flematti *et al.*, 2015) or the as yet unidentified endogenous ligands of the KAI2 (KARRIKIN INSENSITIVE) karrikins receptor (KAI2 ligand, KL, Sun *et al.*, 2016).

Many elements of the KAR/KL perception pathway have been elucidated and are either shared or related to components of the SL perception pathway. The current model suggests that KARs and KLs are perceived by binding the α/β-fold hydrolase KAI2/D14-like protein (Waters *et al.*, 2012; Bythell-Douglas *et al.*, 2013), while SLs bind a related α/β-fold hydrolase called D14 (Hamiaux *et al.*, 2012; Chevalier *et al.*, 2014; Yao *et al.*, 2016). D14 can form a complex with MAX2 (MORE AXILLARY GROWTH2), a leucine-rich repeat F-box protein (Zhao *et al.*, 2015; Yao *et al.*, 2016). It has been hypothesised that KAI2 can also form such a complex with MAX2 though no physical interaction has been confirmed yet (Conn & Nelson, 2016). In addition, the KAR-dependent degradation of KAI2 can occur independently from MAX2, independently of ubiquitination or the activity of the 26S proteasome (Waters *et al.*, 2015a). More recently heat-shock related proteins have been identified as degradation targets of MAX2 in rice (D53, Jiang *et al.*, 2013; Zhou *et al.*, 2013) and *Arabidopsis* (SMXL, SUPPRESSOR OF MAX2-1-LIKE, (Stanga *et al.*, 2013; Soundappan *et al.*, 2015). Thus far, a dichotomy has been proposed with SMAX1 (SUPPRESSOR OF MAX2-1) suppressing karrikin-related *max2* phenotypes (*e.g*., germination and hypocotyl elongation) while other members of the SMXL family, namely SMXL6, SMXL7 and SMXL8, suppress SL-related phenotypes (*e.g*., shoot branching and lateral root density, Waters *et al.*, 2017). While some specificity of SL or KAR/KL signalling is established through the receptors, additional specificity is reinforced through the degradation targets. And, these have been described not merely as suppressors of signalling but also as growth regulators, the activities of which are modulated through SL or KAR/KL signalling (Jiang *et al.*, 2013).

Here a role for KAI2-dependent and MAX2-dependent signalling in regulating specific root growth patterns (skewing, and waving) is demonstrated for the first time, which challenges the current model of SL/KL specificity with regards to their interacting partners from the SMXLs family. KAR_2_ or GR24_rac_ (a synthetic analogue for SL and KAR) are shown to be poor analogues for KLs with regards to root skewing regulation. We propose that KAI2 and MAX2 regulate these growth patterns through a mechanism involving root radial expansion as the mutants’ gravitropic and mechano-sensing responses remained largely unaffected. In addition, this work establishes new connections between MAX2 and SKS3 (*SKU5* Similar), as we show genetic data placing both protein in the same genetic pathway regulating root skewing.

## Materials and Methods

### Plant material and growth conditions

Wild type *Arabidopsis* seeds Columbia-0 (Col) and Landsberg *erecta* (L*er*) were the parental backgrounds for the mutants tested. Seeds for *max2* (*max2-1*) and *Atd14 (Atd14-1,* (Waters *et al.*, 2012) were provided by Prof. Dame Ottoline Leyser (SLCU, Stimberg *et al.*, 2002). Seeds for *max2-7, max2-8, kai2-1, kai2-2, dlk2-1, dlk2-2, dlk2-3,* and KAI2:KAI2 (*kai2-2*) were a gift from Dr. Mark Waters (University of Western Australia, Waters *et al.*, 2012; 2015b). Seeds were surface sterilized by treatment with 70% (v/v) ethanol, followed by a rinse with sterile distilled water then incubation in 10% (v/v) sodium hypochlorite, 0.05% (v/v) Triton X-100 for 5 minutes at 20°C with shaking (1,250 rpm). After a further five washes with sterile distilled water, seeds were placed on the surface of 0.8% (w/v) agar (BD, UK) supplemented with 1/2 MS (Murashige and Skoog including vitamins, pH 5.6, Duchefa, The Netherlands). *Arabidopsis* seeds were stratified in the dark for 2 days at 4°C, before transfer to a growth cabinet under controlled conditions at 23°C, 16h light: 8h dark, and 80 μmol m^-2^ s^-1^ irradiance. Growth plates were vertical unless stated otherwise.

### Root skewing assay

After 9 days, images were taken by scanning plates from the back (*i.e.*, roots were imaged through the agar) using a flat-bed scanner (300 dpi) and root skewing angles were measured in ImageJ (Schneider *et al.*, 2012) using the angle tool. NeuronJ (Meijering *et al.*, 2004) was used to record the *x* and *y* coordinates of the root tips and a marked section of the root. These coordinates were then used to calculate the horizontal growth index (HGI) and vertical growth index (VGI) as previously described (Grabov *et al.*, 2004; Vaughn & Masson, 2011). Waviness was measured as the ratio of the cord to the root length (Grabov *et al.*, 2004; Vaughn & Masson, 2011).

### GR24_rac_ and KAR_2_

Plants were grown for 6 days on the surface of control medium (0.8% (w/v) agar supplemented with 1/2 MS, including vitamins, pH 5.6), then transferred to medium containing racemic GR24_rac_ (LeadGen Labs, USA) or KAR_2_ (Toronto Research Chemicals, Canada) or only the carrier for the test compound as a control (sterile distilled water for KAR_2_ and 0.02% (v/v) acetone for GR24_rac_). Plants were then grown for a further three days before scanning.

### Cell file rotation and root diameter analysis

Images of the root tips from plants grown vertically for 6 days, then placed at a 45° angle from the vertical for a further 3 days, were taken using a Leica DFC365FX camera attached to a Leica M205FA Stereo microscope (Leica Microsystems Ltd, UK) with a Planapo x1.6 objective set to magnification of x80.5. Images were stitched using the LAS X software platform (Leica Microsystems Ltd, UK). Following Wang *et al.* (Wang *et al.*, 2011) cell file rotation (CFR) was defined as the number of epidermal cell files that crossed a 1 mm long straight line drawn down the longitudinal axis of the root from 1.5 to 2.5 mm from the root apex. Using the same images as for CFR measurements, root diameter was measured approximately 2 mm from the root apex using ImageJ (Schneider *et al.*, 2012), three measurements were done per individual root.

### Mechanical stimulation assays for transcriptional response

Plants were grown vertically on the surface of control plates for 9 days were transferred to a sterile buffer solution (0.1 mM KCl, 10 mM CaCl_2_ and 2 mM bis-Tris propane, pH 5.8 adjusted with 0.5 M MES). A total of 30 to 40 seedlings per genotype were transferred into a Petri dish (3 cm in diameter), containing 3 mL of buffer solution, and left to acclimatize on the bench for 3 hours with additional light (15W/865 Lumilux Daylight, maximum intensity: 86 μmol m^-2^ s^-1^). Mechanical stimulation was applied by shaking vigorously for 30 seconds, while control plants remained on the bench. Plants were then left untouched for a further 30 minutes after stimulation before being immersed in RNALater (Sigma Aldrich) for sample collection as described previously. For both assays, RNA was extracted from roots using the RNeasy Plant Mini kit (Qiagen) per manufacturer’s instructions, including an additional DNase digestion step. A LiCl precipitation step was used to purify and concentrate the RNA before downstream qPCR analysis.

### cDNA synthesis and transcript abundance measurement

Complementary DNA (cDNA) was synthesized from 500 ng RNA using the RT QuantiTect reverse transcription kit (Qiagen), following manufacturer’s instructions except that incubation time was lengthened for the gDNA Wipeout step (3 minutes at 42°C) and the cDNA synthesis (25 minutes at 42°C). cDNA was used as template in a quantitative real-time PCR using the SYBR GREEN PCR kit (Qiagen) and the Rotor-Gene 3000 thermocycler (Qiagen) to determine transcript abundance of the genes of interest *Calmodulin-like* (*CML) 12* and *CML24*. qPCR amplification cycle consisted of 5 min at 95°C followed by 40 cycles of 5 s at 95°C and 10 s at 60°C. Melting curves (ramping from 55°C to 95°C rising 1°C each step, with a 5s delay between steps) were checked for unspecific amplification. qPCR traces were analysed using the R qpcR package (relevant parameters: data were normalized and the background subtracted; starting fit model: l4; efficiency estimation: cpD2; refmean: True; baseline subtraction using the average of the first 5 cycles; (Ritz & Spiess, 2008) R package version 1.4-0. 2015) to calculate Ct values. Efficiencies (all > 92%) were calculated using the calibration curve method. For each gene, the expression was calculated following the formula E= (eff^-Ct^). Expression of the genes of interest was normalised against two housekeeping genes *Ubiquitin 10* (*UBQ10*) and *Tubulin 4* (*TUB4*), as followed R_Gene of Interest_ = E_Gene of_ Interest/(sqrt(E_UBQ10_* E_TUB4_)). qPCR primers are listed in Table S1.

### Measurements of cytosolic Ca^2+^ concentration ([Ca^2+^]_cyt_) in response to mechanical stimulation

Col and *max2* (transformed using floral dip with *Agrobacterium tumefaciens* to express (apo)aequorin under a 35S promoter, Dodd *et al.*, 2006)) were used at T3 or T4 generation to determine cytosolic free Ca^2+^ concentration ([Ca^2+^]_cyt_). Equivalence of aequorin levels were determined by discharge assay of luminescence (> 4 million luminescence counts for both Col and *max2*). Plants were grown vertically on solid medium for 7-8 days as described above. Excised root tips (1 cm) were placed in the wells (one root per well) of a white 96-well plate (Greiner Bio-One, UK) and incubated in 100 μL of bathing solution (10 μM coelentrazine, Lux Biotechnology, UK), 0.1 mM KCl, 10 mM CaCl_2_ and 2 mM bis-Tris propane, pH 5.8 adjusted with 0.5 M MES) for 2h in the dark, at room temperature. Luminescence was then recorded every second in a plate-reading luminometer (FLUOstar Optima, BMG labtech, Ortenberg, Germany). After 35 s, 100 μL of bathing solution (without coelentrazine) was injected into the well at 200 μL s^-1^ to cause a mechanical stimulus to the root resulting in a sudden increase in luminescence (“touch response”). The signal was monitored for a further 120 s, when a 100 μL of discharge solution (3 M CaCl_2_, in 30% (v/v) ethanol) was delivered to normalize the luminescence data and calculate [Ca^2+^]_cyt_ (Laohavisit *et al.*, 2012). The [Ca^2+^]_cyt_ touch response of Col and *max2* were then compared.

### Root gravitropism assays

*Arabidopsis* plants were grown vertically for 14 days on the surface of control medium. On the day of the experiment, roots were positioned by aligning their root tips so that they could be imaged together. Plates were then placed vertically in the growth incubator but rotated through a 90° angle thus inducing a 90° change in gravitropic orientation. Root tips were imaged using a Raspberry Pi camera module (http://www.raspberrypi.org/). Images were acquired every 10 min for 10 h. Image analysis was conducted using ARTT (Russino *et al.*, 2013) which tracked the root tip growth and gave the tip orientation and displacement as output. Tip orientation was normalised to the displacement to take into account differences in growth rate.

### Data representation and statistical analysis

Root skewing data were represented using beanplots constructed in the R environment (R Core Team, 2012) using the beanplot package (Kampstra, 2014), to show the variability in root skewing angle. Statistical analyses were also conducted in the R environment. Normal distribution of the data and equality of variance were verified using Shapiro and Levene tests (Lawstat package, Gastwirth *et al.*, 2017), respectively. Significant differences amongst genotypes were verified using one-way Analysis of Variance (ANOVA), followed by Tukey HSD. ANOVAs were conducted on rank values as a non-parametric method, when data did not uphold the assumptions of normality and homoscedasticity. All experiments were repeated at least three times.

## Results

### Mutation in *kai2* and *max2* increases root rightward skew

If KL or karrikins were involved in root skewing then insensitive mutants of *Arabidopsis thaliana* would display an aberrant root-skewing phenotype. Vertically-grown *kai2-1* and *kai2-2* mutants showed significantly increased rightward root skewing compared to the L*er* wild type (α, root tip displacement, viewed from the back of the plate: Fig. 1a, b; Tukey HSD, *p* < 0.01). Vertically-grown *max2-7* and *max2-8* mutants also showed a significant increase in rightward root skewing compared to wild type (Fig. 1a, b; Tukey HSD, *p* < 0.01).

**Fig.1.**
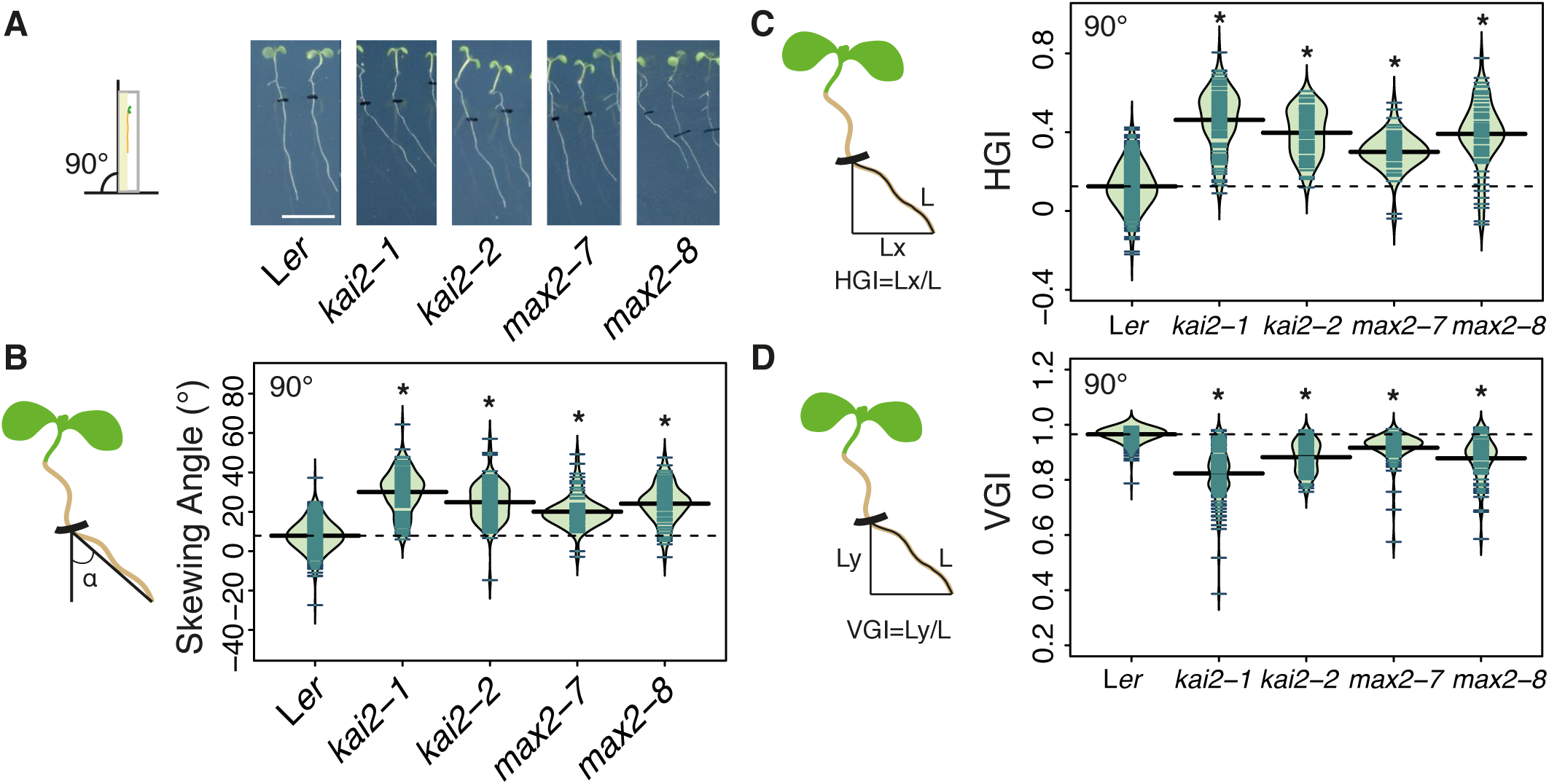
*kai2* and *max2* mutants display an exaggerated rightward root skewing phenotype. A. Seedlings of *kai2-1, kai2-2, max2-7*, and *max2-8*, displayed an exaggerated rightward skew when grown at 90°. The scale bar represents 1 cm. B. The root skewing angle (α) was measured as the deviation from the vertical for plants grown at a 90° angle. C. The increased root skewing of karrikin-insensitive mutants measured as the simple deviation from the vertical could also be noted when measured as an increase in horizontal growth index (HGI) or (D) a decrease in vertical growth index (VGI). Data for each genotype are displayed as a beanplot with the skewing angle of individual roots shown as dark green horizontal lines, while the mean is represented by a thick black horizontal line. The estimated density of the distribution is illustrated by the shaded colour. The dashed line corresponds to the mean for the wild type. Positive values are rightward skews. * indicates significant difference compared to wild type (Tukey HSD, *p* < 0.05). For each genotype, *n* > 65 in 3 separate experiments.

Horizontal Growth Index (HGI; ratio of root tip displacement along the *x* axis to root length, Grabov *et al.*, 2004; Vaughn & Masson, 2011) was also significantly higher in *kai2-1, kai2-2, max2-7*, and *max2-8* compared to wild type (Fig. 1c; Tukey HSD, *p* < 0.01), supporting the skewing angle data and showing increased deviation from vertical by mutant roots. Similarly, the Vertical Growth Index (VGI; ratio of root tip displacement along the *y* axis to root length, Grabov *et al.*, 2004; Vaughn & Masson, 2011) was significantly smaller for *kai2-1, kai2-2, max2-7*, and *max2-8* compared to wild type (Fig. 1b; Tukey HSD, *p* < 0.01). In separate experiments, two complemented *kai2-2* lines (driven by the native promoter KAI2:KAI2 (*kai2-2*), Waters *et al.*, 2015b) showed significantly decreased root skewing angle compared to *kai2-2* (Fig. S1a; Tukey HSD, *p* < 0.05). Overall these data suggest a role for both KAI2 and MAX2 in root skewing.

### *KAI2* and *MAX2* operate through the same genetic pathway

Although no physical interaction has been demonstrated between KAI2 and MAX2, they have been placed in the same signalling pathway through genetic studies of elongated hypocotyl phenotypes (Waters *et al.*, 2012). Here, the double mutant *kai2-2 max2-8* showed a significantly increased rightward root skewing compared to wild type (Fig. 2a, b, Tukey HSD, *p* < 0.05), which was not significantly different from that of *kai2-2* (Fig. 2a, b; Tukey HSD, *n.s.*). That skewing angle of the *kai2-2 max2-8* double mutant was not greater than that of the *kai2-2* single mutant suggests that KAI2 and MAX2 operate in the same pathway. Critically, *d14* mutants that are insensitive to SL but not KAR (Waters *et al.*, 2012) showed no significant increase in root skewing compared to wild type (Fig. 2c, d, ANOVA, *n.s.*). A higher root skewing angle for wild type L*er* plants compared to Col was found, as noted previously (Vaughn & Masson, 2011). Moreover, the root skewing of *dlk2* mutants (Waters *et al.*, 2012) was not significantly different to wild type (Fig. S1b; Tukey HSD, *n.s.*). As the DLK2 protein is related to both KAI2 and D14, overall these data demonstrate a specific role for KAI2 and MAX2 in the regulation of root skewing and thus implicate KL/KAR sensing through these proteins.

**Fig. 2.**
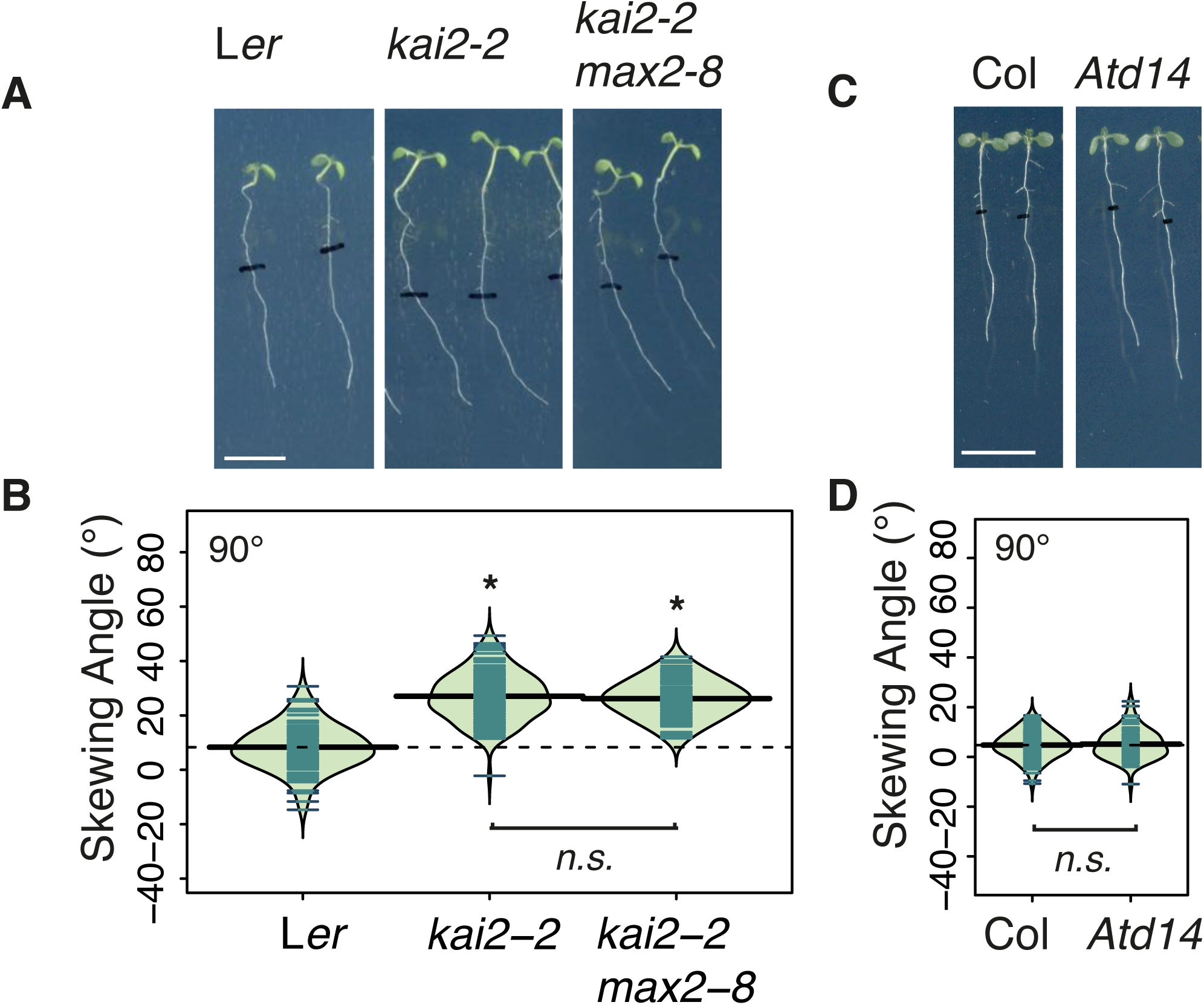
KAI2 and MAX2 regulate root skewing through the same pathway, which does not involve D14. A. Seedlings for the double mutant *kai2-2/max2-8* showed no further increase in root skewing angle compared to *kai2-2* (B). The scale bar indicates 1 cm. Data for each genotype are displayed as a beanplot with the skewing angle of individual roots shown as dark green horizontal lines, while the mean is represented by a thick black horizontal line. The estimated density of the distribution is illustrated by the shaded colour. The dashed line corresponds to the mean for the wild type. * indicates significant difference compared to wild type (Tukey HSD, *p* < 0.05). For each genotype, *n* > 66 in 5 separate experiments. C. Seedlings for the SL-insensitive mutant *Atd14* showed no increased rightward root skewing, and the measured skewing angle was not significantly different from that of the wild type (D). For each genotype, *n* > 73 from 3 experiments.

### KAR_2_ reduces root skewing

In the absence of purified and identified KL compounds, the effect of KAR on root skewing was tested using the potent karrikin KAR_2_ (Nelson *et al.*, 2009; Waters *et al.*, 2015a). The phenotype of *kai2* and *max2* mutants suggests that an impairment in KL perception leads to greater rightward root skewing. Therefore, an increased availability of KL or its analogue KAR_2_ might compensate for a lowered sensitivity of the system and decrease the rightward root skewing. Here there was a significant effect of KAR_2_ in reducing rightward root skewing of L*er* wild type plants with concentrations of 5 and 10 μM (Fig. S2a Tukey HSD, *p* < 0.05). A significant inhibitory effect on primary root elongation of L*er* plants was evident at 10 μM KAR_2_ (Fig. S2b, Tukey HSD, *p* < 0.01).

The presence of 2.5 and 5 μM KAR_2_ in the medium also significantly decreased the root skewing angle of *kai2-2* (Fig. S2b; Tukey HSD, *p* < 0.05). *kai2* plants seem to be more sensitive to KAR_2_ than L*er* plants. The KAI2-independent effect of KAR_2_ on root skewing may also be linked to reduced root elongation, as this was significantly lower in the presence of 5 μM KAR_2_ (Fig. S2b, Tukey HSD, *p* < 0.01) but not at 2.5 ?M (Fig. S2b, Tukey HSD, *n.s.*). It is likely that *kai2* plants are more sensitive to the unspecific toxicity effect of KAR_2_ than L*er* because they lack a mechanism for degradation of KAR and KL. Similarly, the presence of 5 μM KAR_2_ in the medium significantly decreased the root skewing angle of *max2-8* (Fig. S2e; Tukey HSD, *p* < 0.01) as well as primary root elongation (Fig. S2f; Tukey HSD, *p* < 0.01).

### GR24_rac_ has little effect on root skewing

Because of the structural similarities between KAR and the SL analogue GR24_rac_ (Zwanenburg *et al.*, 2009), and the already established role of GR24_rac_ in regulating root growth (Ruyter-Spira *et al.*, 2011; Kapulnik *et al.*, 2011; Rasmussen *et al.*, 2012), we tested the effect of GR24_rac_ on root skewing. A racemic mix of GR24 (GR24_rac_) that can also be perceived by KAI2 (Scaffidi *et al.*, 2014; Waters *et al.*, 2015a) was tested at 1 and 5 μM as greater concentrations tend to have a toxicity effect on root growth (Ruyter-Spira *et al.*, 2011). Treatment with GR24_rac_ led to a small increase in rightward root skewing in L*er* plants at 1 μM (Fig. S3a, Tukey HSD, *p* < 0.01) but not at 5 μM GR24_rac_ (Tukey HSD, *n.s.*). There was no significant effect of 1 or 5 μM GR24_rac_ on *kai2-1* root skewing (Tukey HSD, *n.s.*). There was no significant effect of 1 μM GR24_rac_ on root skewing of *kai2-2* (Tukey HSD, *n.s.*) and there was a small but significant decrease in *kai2-2* root skewing with 5 μM GR24_rac_ (Tukey HSD, *p* < 0.01). There was no significant effect of 1 or 5 μM GR24_rac_ on the root skewing of Col plants (Fig. S3b, ANOVA, F_(2,261)_=1.26, *n.s.*). There was a small but significant increase in root skewing angle for *max2-1* plants in the presence of 1 μM GR24_rac_ (Tukey HSD, *p <* 0.05) but not 5 μM GR24_rac_ (Tukey HSD, *n.s.*), while *d14* plants did not respond to the presence of GR24_rac_ (ANOVA, F_(2,184)_=1.31, *n.s.).* Overall, GR24_rac_ had little effect on root skewing especially in comparison with the root skewing angles of *kai2* and *max2* mutants, and as such is a poor KL analogue with regards to the regulation of root skewing.

### MAX2 regulation of skewing operates through SMXL6,7,8 but not SMAX1

The involvement of MAX2 degradation targets, SMAX1 (SUPPRESSOR OF MAX2-1) and SMXL (SUPPRESSOR OF MAX2-1-LIKE, (Stanga *et al.*, 2013) in regulating root skewing was examined. D53, a homologue of SMXL6,7,8 in rice, forms a complex with D14 and D3, and is degraded following SL treatment (Jiang *et al.*, 2013; Zhou *et al.*, 2013). The current mechanistic model for *Arabidopsis* is that SMAX1 is important for the KL part of the signalling pathway whereas SMXL6,7,8 are more relevant to the SL part of the pathway (Soundappan *et al.*, 2015).

Here we tested the hypothesis that the loss of SMAX1 or SMXL6,7,8 proteins would affect root skewing. Both *smax1-2* and *smxl6,7,8* mutants had a significantly decreased skew compared to wild type (Fig. 3a, b; Tukey HSD, *p* < 0.01), thus suggesting that the abundance of these proteins is important in regulating the skew. The hypothesis that the MAX2-dependent regulation of SMAX1 and SMXL6,7,8 protein abundance is relevant to the root skewing phenotype was then tested. For this, the root skewing angle of the *max2 smax1-2* double mutant as well as the quadruple mutant *max2-1 smxl6,7,8* was measured. If MAX2 were to affect root skewing exclusively through the abundance of SMAX1 or SMXL6,7,8, then the presence of the *max2* mutation should have no effect on the root skewing phenotype of *smax1-2* or *smxl6,7,8*. Here, a significant increase in root skewing angle in the *smax1-2 max2-1* double mutant compared to *smax1* (Tukey HSD, *p* < 0.01) was observed, but there was no further increase in *max2-1 smxl6,7,8* compared to *smxl6,7,8* (Fig. 3b; Tukey HSD, *n.s.*). Thus, we conclude that the regulation of root skewing by MAX2 is dependent on SMXL6,7,8 rather than SMAX1.

**Fig. 3.**
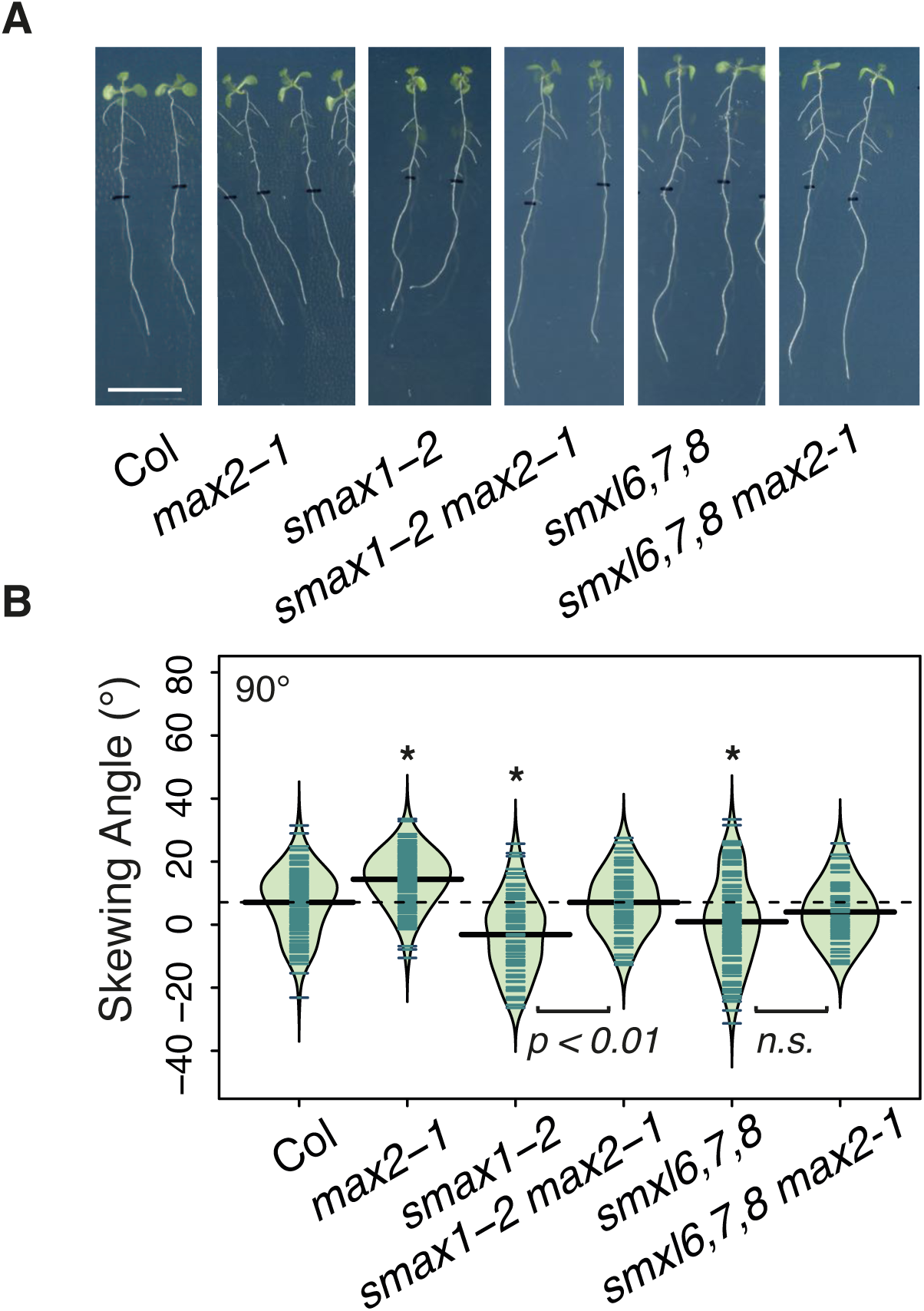
Involvement of SMAX1 and SMXLs in root skewing. A. Seedlings for Col, *max2-1, smax2-1, smax2-1/max2-1, smxl6,7,8* and *smxl6,7,8/max2-1* showing root skewing while grown at 90°. The scale bar represents 1 cm. B. Data for each genotype are displayed as a beanplot with the skewing angle of individual roots shown as dark green horizontal lines, while the mean is represented by a thick black horizontal line. The estimated density of the distribution is illustrated by the shaded colour. The dashed line corresponds to the mean for the wild type. * indicates a significant difference compared to wild type (Tukey HSD, *p* < 0.05). For each genotype, *n* > 58 from 6 experiments.

### KAI2 and MAX2 negatively regulates both skewing on a tilted surface and waving

Positioning plates at a 45° angle rather than 90° increases root skewing angle. A significant increase in rightward root skew angle was observed here for the L*er* wild type grown at a 45°plate angle (Fig. 4a, b, ANOVA, F_(1,510)_=134.9, p < 0.001), whilst *kai2-1, kai2-2, max2-7*, and *max2-8* also showed a significantly increased rightward root skewing angle compared to L*er* (Fig. 3a,b; Tukey HSD, *p* < 0.01). The increase in mutant root skew relative to wild type was maintained at the 45° plate angle compared to growth at 90°, indicating that loss of KAI2 or MAX2 did affect the mutant’s ability to sense and respond to the tilt.

**Fig. 4.**
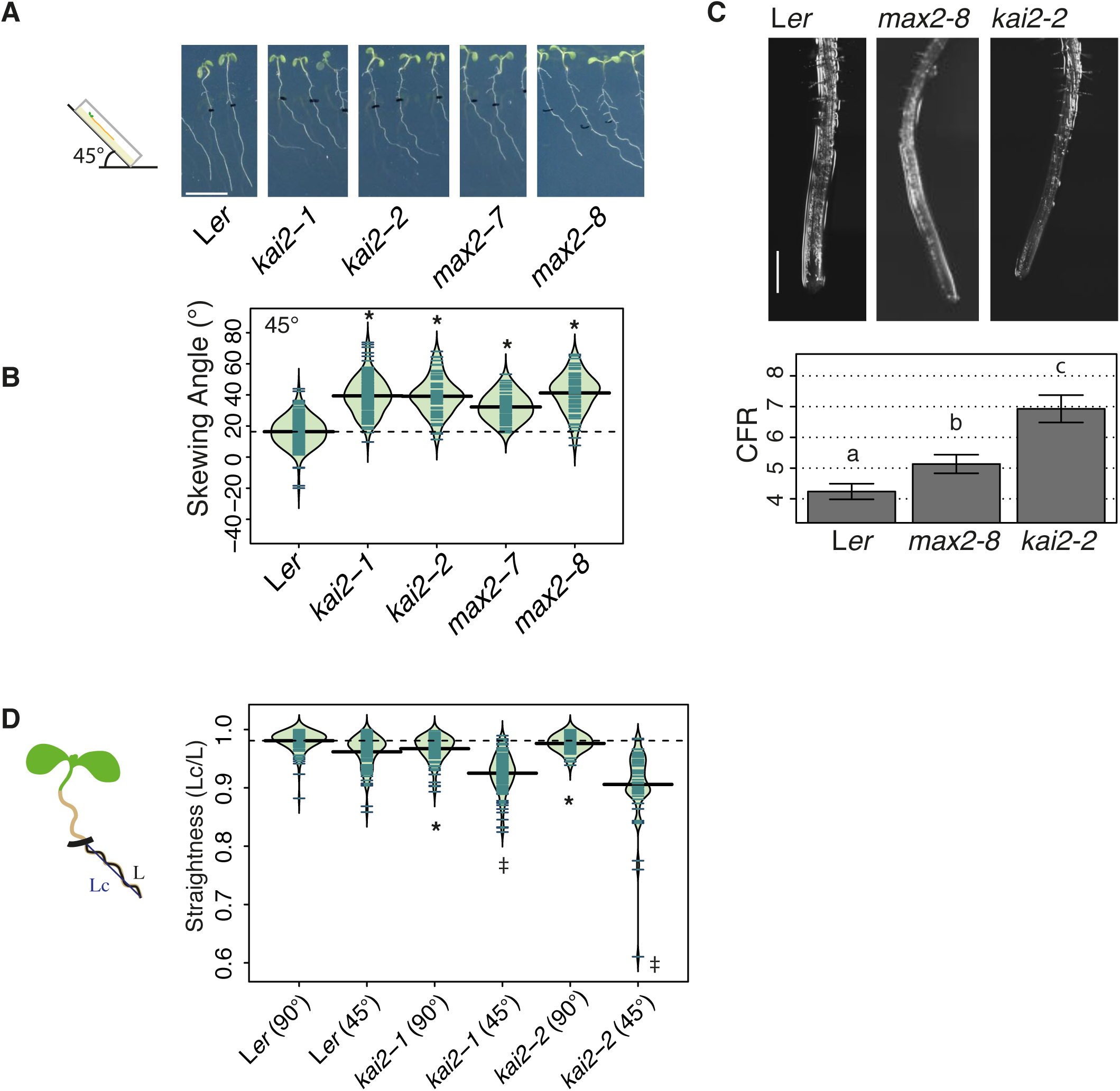
*kai2* and *max2* increased rightward root skewing when placed at 45°. A. Seedlings of *kai2-1, kai2-2, max2-7*, and *max2-8*, displayed an exaggerated rightward skew when grown vertically for six days then placed at 45° for 3 days. The scale bar represents 1cm. B. The root skewing angle (α) was measured as the deviation from the vertical for plants grown at a 45° angle for 3 days. C. Both *max2-8* and *kai2-2* mutants show increased cell file rotation (CFR) indicating that the root epidermal cells were twisting more compared to those of the wild type. CFR was measured as the number of epidermal cells that crossed a 1 mm line 1.5 to 2 mm from the root tip. Plants were grown at 45°. Data shown as mean ± SE, *n* = 28-42 plants obtained in 4 separate experiments. Letters indicate significant differences (Tukey HSD, *p* < 0.05, except for the comparison L*er*-*max2-7* where the difference was significant at the 10% limit, pairwise *t*-test, *p* < 0.05). The scale bar indicates 500μm. D. The straightness (measured as the ratio of the chord Lc to root length L; Grabov *et al.*, 2004; Vaughn & Masson, 2011) of seedling roots from wild type, *kai2-1* and *kai2-2* decreased when plants were grown at 45° compared to 90° (shown in brackets behind genotype). Data for each genotype are displayed as a beanplot with the straightness of individual roots shown as dark green horizontal lines, while the mean is represented by a thick black horizontal line. The estimated density of the distribution is illustrated by the shaded colour. The dashed line corresponds to the mean for the wild type. * indicates significant difference at the 5% level compared to wild type grown at 90°, while ‡ indicates a significant difference to wild type grown at 45°. For each genotype, *n*58 in 3 separate experiments.

Although mechanistic models for root skewing vary (Roy & Bassham, 2014), the rotation of epidermal cell files is considered to be an important feature (Sedbrook *et al.*, 2002; Oliva & Dunand, 2007; Wang *et al.*, 2011). Right-handed cell file rotation was significantly increased in both *kai2-2* (mean ± SEM 6.93 ± 0.44 cell mm^-1^; Tukey HSD, *p* < 0.01) and *max2-8* (5.13 ± 0.30 cell mm^-1^; Tukey HSD, *p* = 0.08) compared to L*er* wild type (4.24 ± 0.25 cell mm^-1^; Fig. 3c).

Increased root skewing is often also accompanied by increased root waving (Roy & Bassham, 2014) - a decrease in root straightness calculated as the ratio of the cord over the root length (*i.e.*, straight roots have a ratio of 1 and the lower the ratio the less straight/more wavy the root; Grabov *et al.*, 2004; Vaughn & Masson, 2011). Growth on a tilted surface can also decrease straightness (Roy & Bassham, 2014). When grown at 90° plate angle, both *kai2-1* and *kai2-2* showed a significantly decreased straightness compared to L*er* wild type (Fig. 4d; Tukey HSD, *p* < 0.05) and similarly when grown at 45° (Fig. 4d; Tukey HSD, *p* < 0.01). L*er* was significantly less straight at 45° compared to 90° (Tukey HSD, *p* < 0.01). These data show that KAI2 is involved in the negative control of both skewing and waving when plants are grown vertically and at an angle.

### *kai2* and *max2* can support a normal mechano-sensing transcriptional response

The growth responses of the *kai2* mutants on tilted plates suggested that the mutation does not affect the root tip’s ability to sense the increased mechanical impedance afforded by the inclined growth medium. Rather, that the *kai2* mutants have an exaggerated root skew when grown on a tilted surface suggests that downstream responses are impaired. To test for a role for KAI2 in mechano-sensing responses seedlings were subjected to mechanical stress prior to determination of root transcript levels of *CML12* and *CML24* (CALMODULIN LIKE PROTEIN,Fig. 5a). These transcripts are known to increase upon mechanical stimulation (Braam & Davis, 1990). These tests also addressed *max2* and *d14* in the Col background (Fig. 5b). Mechanical stimulation caused significant upregulation of *CML12* and *CML24* transcript in roots of all genotypes tested (ANOVA, *p* < 0.001) but no mutants responded significantly differently to their wild type. Thus, the data suggest that root transcriptional mechano-responsiveness is not drastically altered in either KL- or SL-insensitive mutants.

**Fig. 5.**
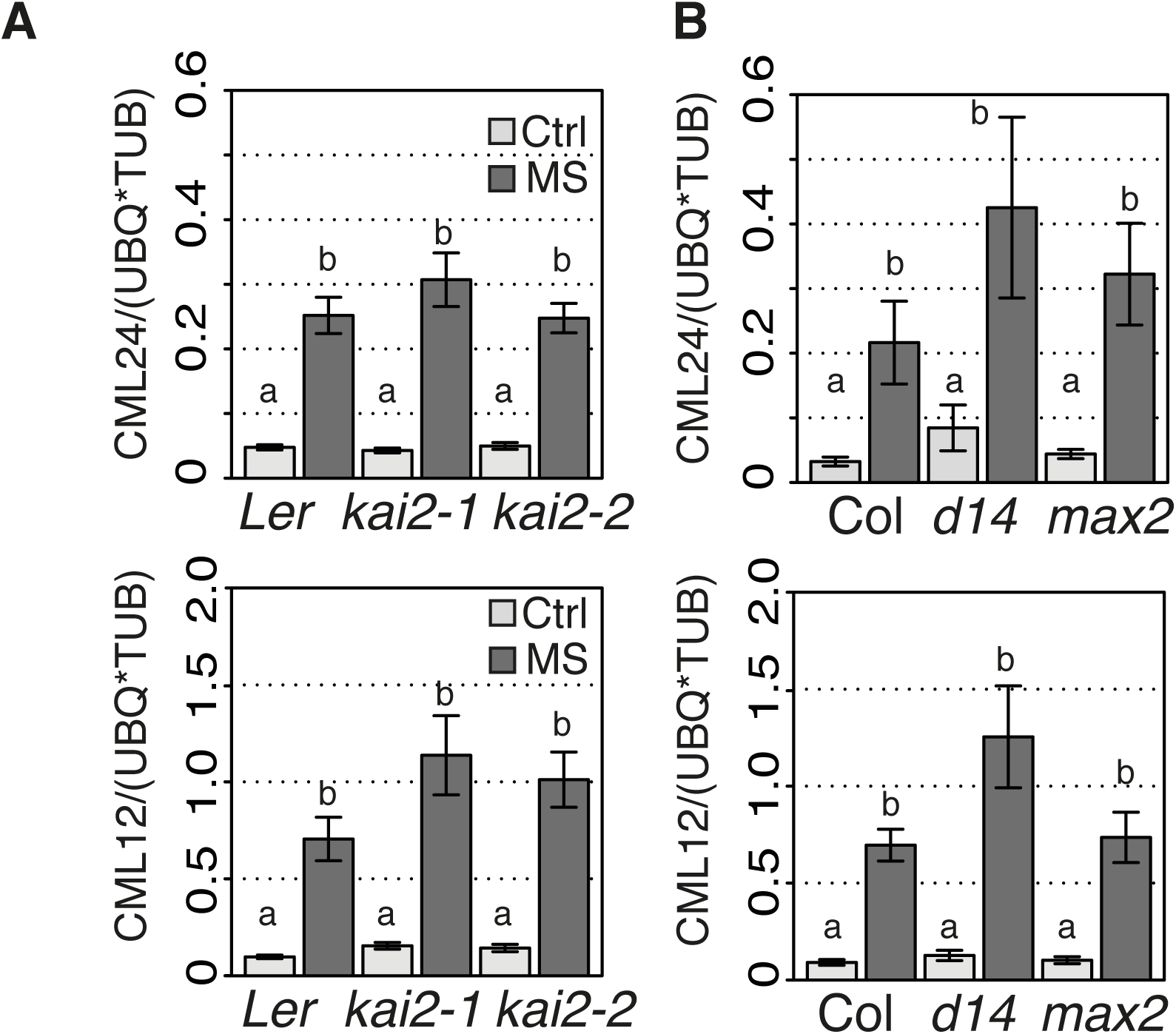
*kai2, max2, d14* mutants support normal transcriptional response to mechano-stimulus. Karrikin- and SL-insensitive mutants showed a normal response to mechanical stimulation at the transcript level. Nine day-old seedlings of wild type and mutants *kai2-1* and *kai2-2* (A), and *d14, max2-1* (B) were mechanically stimulated (MS) for 30 seconds, then collected 30 min later for transcript analysis of touch-sensitive genes *CML12* and *CML24,* relative to housekeeping genes *Tubulin 4* and *Ubiquitin 10*. The means of 6-9 replicates from 3 independent experiments are shown, each replicate based on the RNA extracted from roots of 30 to 40 seedlings. Data are shown as mean ± SE, letters indicate significant differences (Tukey HSD, *p* < 0.05).

As a final test for alteration in mechano-sensing and response, *max2* (as the common lesion in KL- and SL-pathways) was transformed to express (apo)aequorin as a reporter of cytosolic free Ca^2+^ ([Ca^2+^]_cyt_). [Ca^2+^]_cyt_ increases transiently in response to mechano stimulation, acting as a second messenger (Knight *et al.*, 1991). There was no significant difference between baseline level pre-injection and post-injection for Col (t-test, *n.s.*) or *max2* (t-test, *n.s.*). There was no significant difference in the amplitude of the touch-induced peak increase in [Ca^2+^]_cyt_ between genotypes (Fig. 6; *t*-test, *n.s.*). However, the total Ca^2+^ mobilised over the recording period (excluding the discharge) for *max2* (33.99 μMs ± 0.57) was significantly higher than that for Col (29.91 μMs ±0.49; *t*-test, p<0.01).

**Fig. 6.**
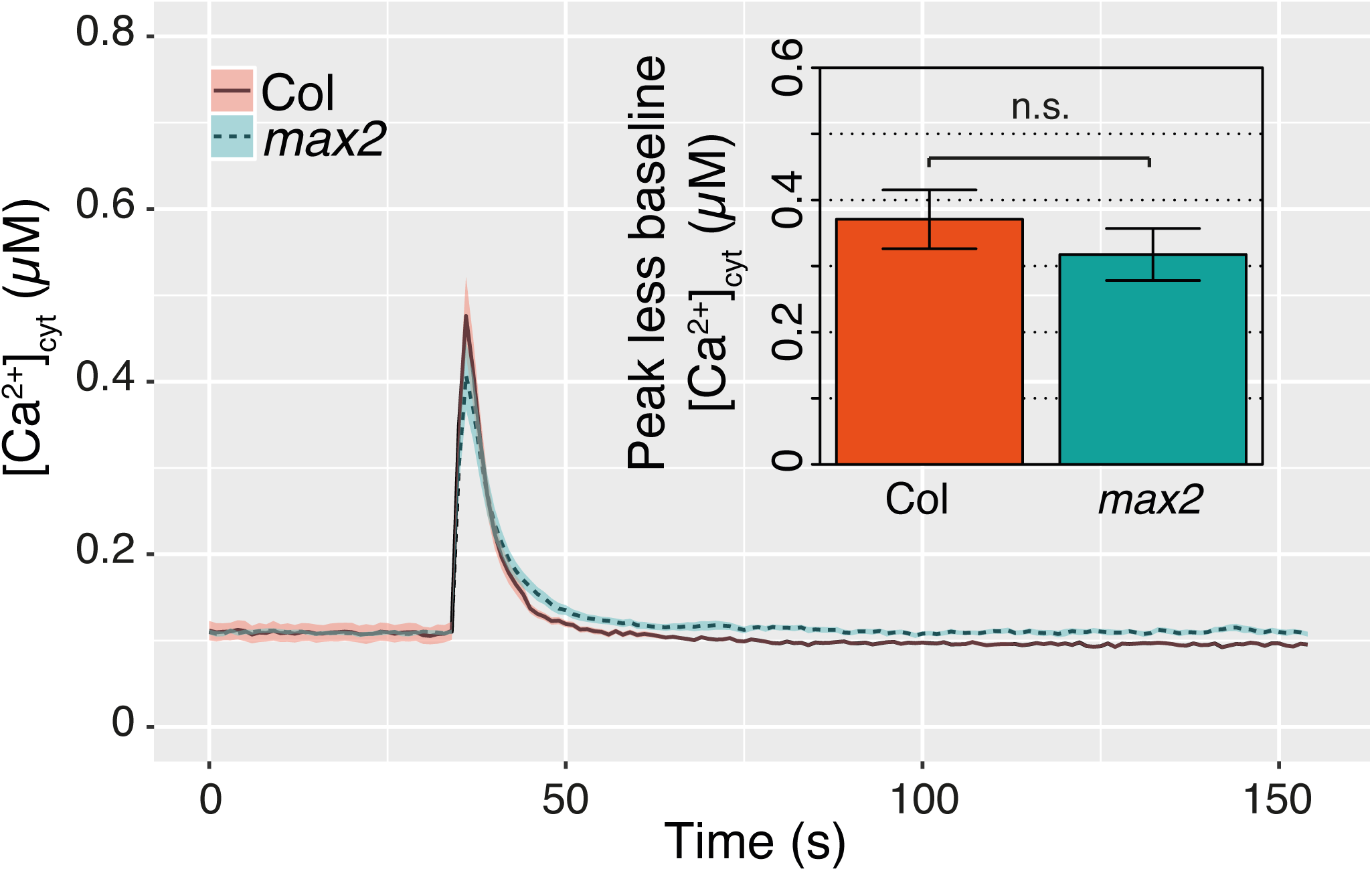
Mechano-stimulated [Ca^2+^]_cyt_ increase in *max2* root tips. Individual excised root tips of Col and *max2* expressing (apo)aequorin as a [Ca^2+^]_cyt_ reporter were mechanically stimulated by addition of buffer at 35s. The mean ± SE of 40 to 67 roots in 5 independent trials are shown. Inset: Mean ± SE maximal [Ca^2+^]_cyt_ increment in response to stimulus (peak response minus baseline).

### *kai2* has a slower early gravitropic response

Agravitropic mutants can also show an increased root skewing (Okada & Shimura, 1990). To investigate whether an aberrant gravitropic response of *kai2-2* plants contributed to their skewing phenotype, root tip orientation was monitored every 10 min after gravistimulation for 10 h. Both *kai2-2* and wild type responded significantly with a change of tip orientation over time (Fig. 7; ANOVA F_(1,4022)_ = 46.8, *p* < 0.001). Comparisons of the responses (normalised for elongation rate) using ANOVA showed that there was a significant interaction between time and genotype (ANOVA, F_(1, 4022)_ = 40.9, *p* < 0.01), indicating a difference in gravitropic response between genotypes. *kai2-2* root tip angle started to decrease later than L*er*. After 100 min, the angle of *kai2-2* was significantly higher than that of L*er* (ANOVA, F_(1,64)_= 4.4, *p* < 0.01) but at 600 min there was no significant difference (ANOVA, F_(1,64)_=0.24, *n.s.*). Overall, the difference in gravitropic response between *kai2-2* and L*er* may be a small contributory factor to root skewing, but occurring only in the early stages of the response.

**Fig. 7.**
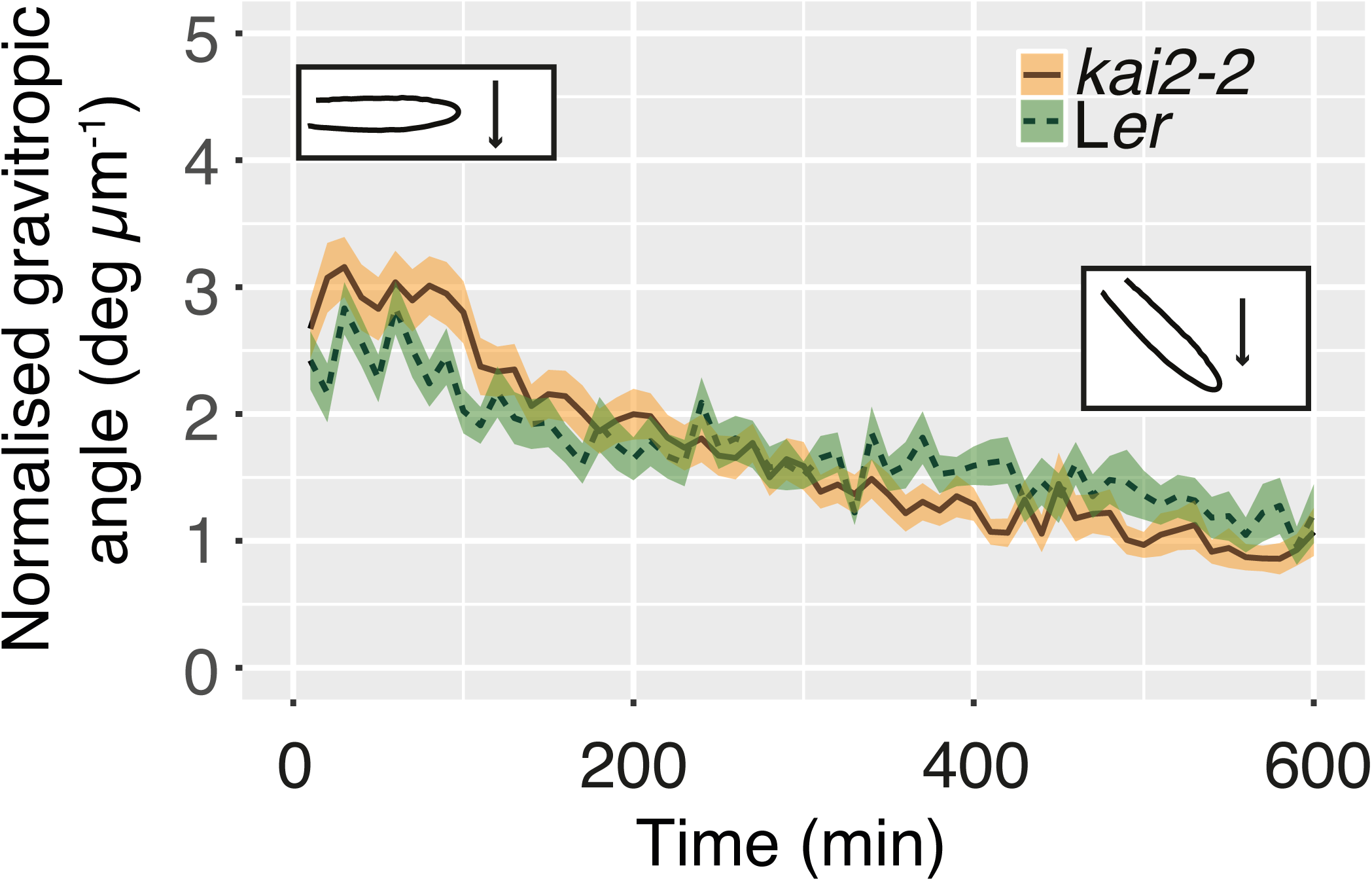
Gravitropic response of *kai2* is slower than wild type’s. The tip orientation of roots from wild type and *kai2-2* was recorded every 10 min and for 10 h after a change in gravitropic orientation. The change in tip orientation was normalised to the tip displacement to take into account differences in growth rate between genotype. Data are shown as mean ± SE, *n* = 16-22 plants obtained in 5 experiments.

**Fig. 8.**
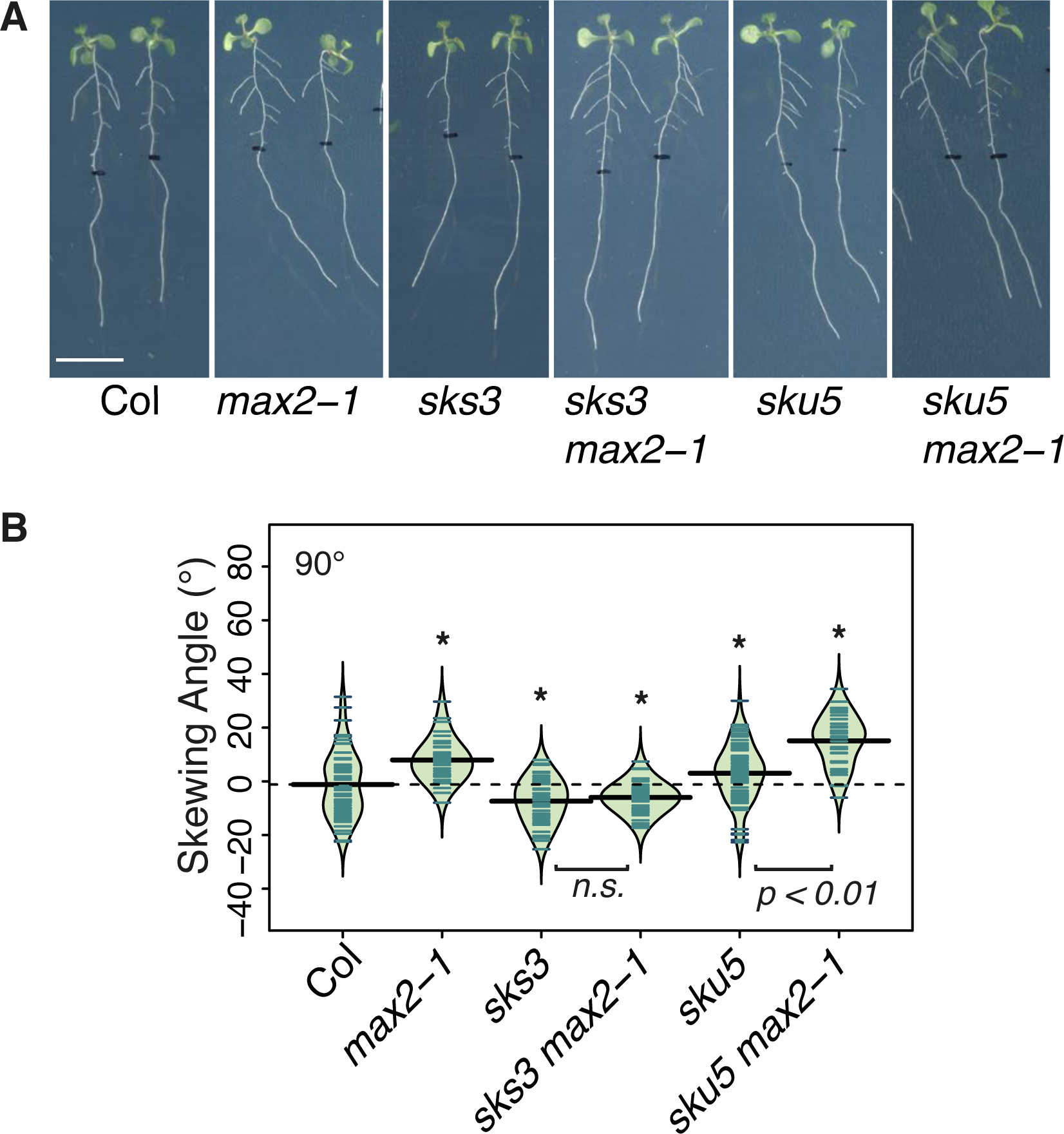
MAX2 regulation of root skewing involves SKS3 and SKU5. A. Seedlings of Col, *max2-1, sks3, sks3/max2-1, sku5, sku5/max2-1* mutants grown at 90°. The scale bar represents 1 cm. B. Data for each genotype are displayed as a beanplot with the skewing angle of individual roots shown as dark green horizontal lines, while the mean is represented by a thick black horizontal line. The estimated density of the distribution is illustrated by the shaded colour. The dashed line corresponds to the mean for the wild type. * indicates a significant difference compared to wild type (Tukey HSD, *p* < 0.05). For each genotype, *n* > 34 in 3 separate experiments.

### MAX2 regulation of root skewing involves SKS3 and SKU5

Similarly to the *kai2* and *max2* mutants, mutant plants deficient in the SKU5 protein that is linked to the plasma membrane by a glycosylphosphatidylinositol (GPI) anchor also showed an increased rightward root skewing phenotype, increased CFR with no change in gravitropic response (Sedbrook *et al.*, 2002). In our experiments, *sku5* also displayed a rightward skew when grown vertically that was significantly greater than the wild type (Fig 8b, Tukey HSD, *p* < 0.05). There was no further increase in *sku5 max2* compared to *max2* (Tukey HSD, *n.s.*), showing that SKU5 and MAX2 can regulate root skewing in the same pathway but as the skewing angle of the *sku5 max2* mutant was significantly higher than that of *sku5* (Tukey HSD, *p* < 0.001) this suggests that part of the MAX2 pathway is SKU5-independent. The *sks3* (*sku5 similar 3*) mutant deficient in a SKU5-related protein also showed a decreased rightward root skewing (Tukey HSD, *p* < 0.05) that was maintained even in the absence of MAX2 (comparison *sks3*: *sks3 max2-1*, Tukey HSD, *n.s.*). These data suggest that the abundance of SKS3 protein may itself regulate root skewing and that the abundance of this protein may be regulated through the MAX2 pathway. *sks3* and *sku5* do suppress the high LRD of *max2* mutants (Fig. S4) as well as the decreased germination rate (Fig. S5). Thus, our data suggest that members of the SKU/SKS at least SKS3 are degradation targets for the MAX2 pathway, and in the case of SKS3 specifically regulating of root skewing. The genetic link established here between MAX2 and SKU/SKS family points towards a role of MAX2 in regulating, through SKS3, a cell wall-dependent process.

### KAI2 and MAX2 positively regulate root diameter

Given the subtle responses in terms of gravitropism and mechanical stimulation versus the clear increase in CFR and link with members of the SKU/SKS family, we hypothesise that in both the *kai2* and *max2* mutants the root skewing phenotype arises due to a restriction of cell growth. This is supported by our finding that the mean root diameter of the mutants was significantly narrower than that of wild type (Fig. S6, L*er* 166.43 μm ± 1.79, *kai2-2* 155.57 μm ± 1.41, *max2-8* 146.59 μm ± 1.67; Tukey HSD *p* < 0.001), suggesting that root radial expansion may be restricted.

## Discussion

Evidence here demonstrating a role for KAI2 and MAX2 in regulating root skewing and waving in *Arabidopsis* reinforces the idea that plant endogenous KL can act as a phytohormone (Conn & Nelson, 2016). This is the first root growth phenotype characterised for karrikin-insensitive mutants in a non-host species (Gutjahr *et al.* 2015).

### KAI2 and MAX2 as new regulatory components for root skewing

The characterization of different root skewing and waving abilities amongst *Arabidopsis* ecotypes strongly suggests that the surface-dependent growth patterns represent an adaptive response relevant to natural soil conditions (Vaughn & Masson, 2011; Schultz *et al.*, 2017). Mutants have proved useful in identifying new components of the machinery regulating root skewing in *Arabidopsis*. Here the increased root skewing phenotype of *kai2* and *max2* suggests that both KAI2 and MAX2 negatively regulate root skewing. Since these two proteins are involved in the perception of KAR and KL this provides evidence supporting a role for KL in regulating root skewing. Previous studies have shown an involvement of the SL pathway in the regulation of root system architecture (although skewing and waving were not reported) (Ruyter-Spira *et al.*, 2011; Kapulnik *et al.*, 2011; Rasmussen *et al.*, 2012). We found no evidence supporting a role for endogenous SL in root skewing, since the *d14* mutant impaired in the perception of SL does not show a root skewing phenotype.

For phenotypes such as elongated hypocotyls or increased seed dormancy (Waters *et al.*, 2012), KAR_2_ acts as a good synthetic analogue for KL (Conn & Nelson, 2016). However, this is not the case for root skewing, in which a high concentration of KAR_2_ is necessary to induce a phenotype and is KAI2 independent. A similarly high (10 μM) concentration of the less potent KAR_1_ could also induce a KAI2-dependent reduction in hypocotyl length (Waters *et al.*, 2012). KAR_1_ is also less potent than KAR_2_ in targeting the degradation of KAI2 protein (Waters *et al.*, 2015a). Results here suggest that KL represent a family of related compounds that can regulate different aspects of plant development, and that the KL responsible for regulating root skewing may differ from the KL responsible for regulating hypocotyl elongation and seed germination. Many SL compounds have been purified thus far (Bouwmeester *et al.*, 2007), perhaps structural diversity is also the case for KL. Although KAR_2_ can regulate root skewing, the high concentration required plus the independence from KAI2 and MAX2 suggest that KAR_2_ is likely non-specific and does not represent a good analogue of KL.

### Root skewing phenotype suggests new links between MAX2 and SKS proteins

The mechanism by which KAI2 and MAX2 regulates root skewing remains elusive but must involve a differential growth leading to increased epidermal cell file rotation. We found no evidence supporting a role for KAI2 and MAX2 in regulating the root mechano-sensing transcriptional response and only a very subtle effect of MAX2 on mechano-stimulated [Ca^2+^]_cyt_ response. The link established here between MAX2 and SKU5 as well as MAX2 and SKS3 suggests the possibility that the *max2* skewing phenotype is linked to cell wall modification or integrity. Amongst the 11 highly probable skew gene candidates identified in *Arabidopsis* roots using microarrays, three were associated to the cell wall either because of their physical location (*PAP24*), or because of their role in cell wall integrity (*DIN2*) or formation (*MIOX4*, Schultz *et al.*, 2017). *SKS15* also presented an expression pattern indicative of a possible role in root skewing in this study. However, analysis of the cell wall composition using Fourier transform infrared spectroscopy and analysis of neutral sugars revealed no differences between *sku5* mutant and wild type (Sedbrook *et al.*, 2002).

Several lines of evidence suggest that KL and KAR affect cell wall composition. First, amongst the 133 genes that are differentially regulated 24h post-imbibition with 1 μM KAR_1_, 11 relate to cell wall (Nelson *et al.*, 2010). Genes annotated as being part of the ‘plant-cell type cell wall’ category of the GO cellular component were significantly enriched in the set of genes regulated by KAR_1_. Amongst those genes, *sks17* (*SKU5 similar 17*) was found to be upregulated 2.2-fold upon treatment with KAR_1_. It is unclear whether the levels of proteins also increase upon treatment with KAR_1_. Second, metabolomic analyses showed reduced levels of flavonoids contributing to lignin composition (including ρ-coumaric acids and ferulic acids) in *max2* roots compared to wild type roots under control conditions (Walton *et al.*, 2016). These are also good indicators of lower levels of cutin monomer, which signals in the AM-root symbiosis (Wang *et al.* 2012). Thus, an altered cell wall would fit with the impairment in the early events leading to the establishment of KAI2-dependent AM symbiosis in host species (Gutjahr *et al.*, 2015) and could feasibly influence root skewing and waving.

### Root skewing phenotype challenges the current model for the SMXLs

Soundappan *et al.* (2015) suggested specific relationships between SMAX1 and KAI2-KAR/ KL-regulated growth and between SMXL6,7,8 and D14-SL-regulated growth. However, data here do not support the idea that there is a clear dichotomy in terms of the degradation-target proteins involved in the perception pathways for SL and KL. Rather the data support a role for MAX2 in regulating the skew in a D14-independent pathway through SMXL6,7,8 rather than SMAX1. However, this is complicated by the fact that SMAX1 itself appears to regulate root skewing. One explanation for this observation might be that SMAX1 regulates the skew indirectly via the regulation of SMX6,7,8. In this scenario, the *smax1-2* mutant has a skewing phenotype because of a decreased level of SMXL6,7,8, proteins. The lack of direct effect of SMAX1 on root skewing is also supported by the fact that there is no further increase in root skewing in the *smxl6,7,8 max2* mutant compared to *smxl6*,7,8. In the quadruple mutant SMAX1 protein levels should be different because SMAX1 is regulated through MAX2 (Stanga *et al.*, 2013; Soundappan *et al.*, 2015). Similarly, the level of SMXL6,7,8 should be higher in the *smax1-2 max2-1* compared to *smax1-2*, thus leading to the observed increase in root skewing and supporting a role for SMXL6,7,8 in regulating root skewing.

In addition, a role was found for KAI2 and MAX2 but not D14 in regulating root skewing. Overall, this suggests that with regards to the regulation of root growth patterns, SMXL6,7,8 as well as SMAX1 may be involved in the MAX2-dependent regulation of skewing, which was also found to be KL-dependent rather than SL-dependent. Much may depend on the spatial localisation of proteins. SMAX1 is expressed in the root cap, while SMXL6, 7 and 8 are also present in the vasculature or mature roots (Soundappan *et al.*, 2015). *KAI2* expression could be found preferentially in the vasculature (Brady *et al.*, 2007) potentially favouring interaction with SMXL6, 7 or 8.

## Conclusions

Root positioning in the soil is critical in terms of regulating access to nutrients and water, but also interaction with neighbours (Fang *et al.*, 2013). The regulation of root positioning is dependent on both the genetic and environmental response. While it is difficult to argue for the field-relevance of root skewing patterns observed on the surface of agar plates, the characterization of different root skewing and waving abilities amongst *Arabidopsis* ecotypes strongly suggests that the surface-dependent growth patterns represent an adaptive response relevant to natural soil conditions (Vaughn & Masson, 2011). The involvement of both KAI2 and MAX2 suggests a role for a potential new phytohormone KL, in regulating root skewing and waving.

## Acknowledgments

We thank Dr. Mark Waters and Prof. Dame Ottoline Leyser for providing seeds and commenting on the manuscript. Thank you also to Prof. David Nelson for providing seeds and Daniel Safka for support in setting up the raspberry Pi system. We are grateful to Dr. Uta Paszkowski, Prof. Alex Webb, Dr. Siobhan Braybrook, Prof. Sidney Shaw and Dr. Jenny Mortimer for interesting discussions. Work was supported by the Broodbank Trust, the Newton Trust, the Gatsby Foundation, and the BBSRC Doctoral Training Programme (BB/J014540/1)

## Authors Contribution

S.M.S. and J.M.D. planned and designed the research. S.M.S., Y.G., E.M. and F.J. performed experiments and analysed data. S.M.S. and J.M.D. wrote the manuscript.

## Conflict of interest

The authors have no conflict of interest to declare.

**Table S1:**
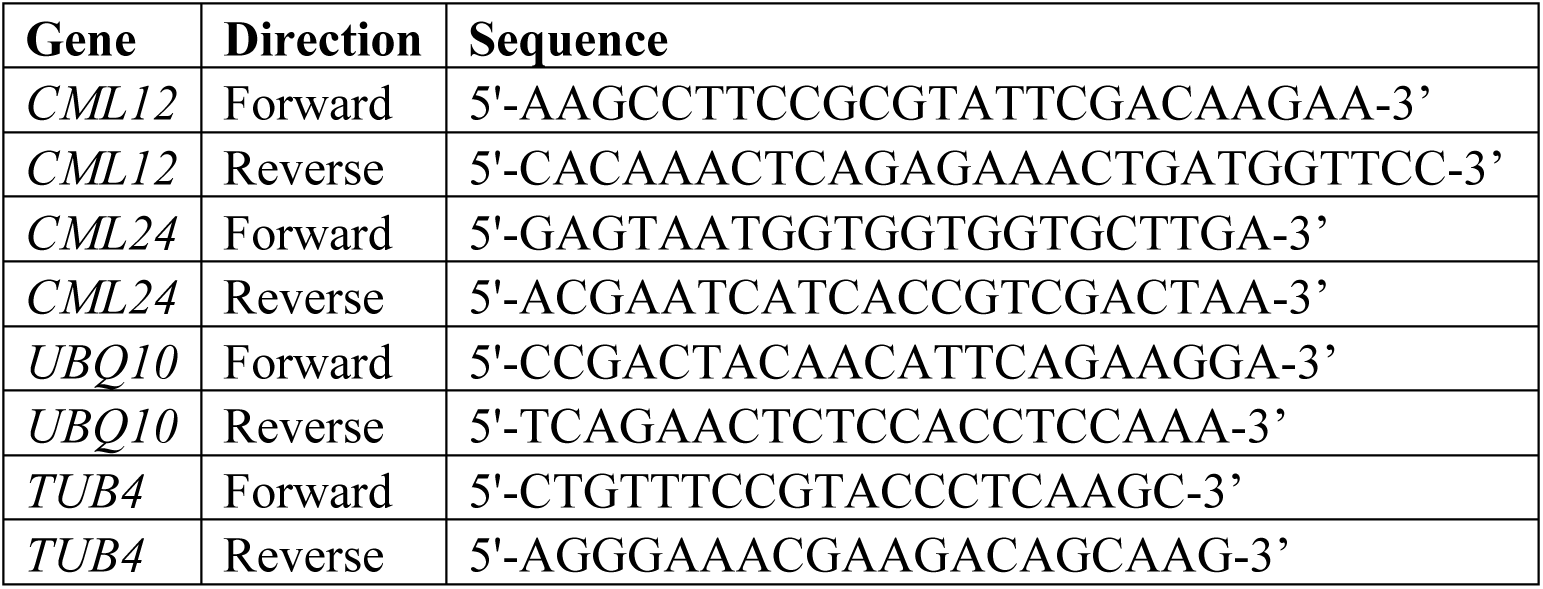
Primer sequences used in qPCR analysis.

## Figure legends

Fig. S1. The root skewing angle of complemented lines of the *kai2-2* mutant was reduced compared to the *kai2-2* mutant but remained higher than that of the L*er* wild type (A). 10G and 12H are *kai2-2* lines complemented by *KAI2* expression under the native promoter. Data for each genotype are displayed as a beanplot with the skewing angle of individual roots shown as dark green horizontal lines, while the mean is represented by a thick black horizontal line. The estimated density of the distribution is illustrated by the shaded colour. The dashed line corresponds to the mean for the wild type. * indicates a significant difference compared to wild type (Tukey HSD, *p* < 0.05) while ‡ indicates a significant difference compared to *kai2-2* (Tukey HSD, *p* < 0.05). For each genotype, *n* > 48 in 4 separate experiments. (B) The root skewing angle of seedlings for three mutant alleles of *dlk2* showed no further increased compared to wild type. There is no significant difference between root skewing angle of *dlk2* alleles and wild type (Tukey HSD, *n.s.*). For each genotype, *n* > 98 in 4 separate experiments.

Fig. S2. Effect of KAR_2_ on root skewing and primary root elongation in *kai2* and *max2*. Root skewing angle of L*er* (A) and *kai2-2* (C) plants grown under control conditions or in the presence of 2.5, 5 or 10 μM KAR_2_ in the medium and, *max2-8* (E) grown under control conditions or in the presence of 5 μM KAR_2_ in the medium. Root elongation over a three-day period when L*er* (B), *kai2-2* (D) and *max2-8* (F) plants were exposed to KAR_2_. Data for each genotype are displayed as a beanplot with the skewing angle of individual roots shown as dark green (or purple for the root elongation data) horizontal lines, while the mean is represented by a thick black horizontal line. The estimated density of the distribution is illustrated by the shaded colour. The dashed line corresponds to the mean for the control conditions. * indicates a significant difference compared to control conditions (Tukey HSD, *p* < 0.05). For each treatment and genotype combination, *n* > 64 (except for *kai2-2* under 2.5 and 10 μM where *n* 30) in at least 3 independent experiments.

Fig. S3. Effect of GR24 on root skewing in *kai2, max2* and *d14*. Root skewing angle of Col (A), *max2-1* (B) and *d14* (C) plants grown under control conditions or in the presence of 1 or 5 μM GR24 in the medium. Data for each genotype are displayed as a beanplot with the skewing angle of individual roots shown as dark green horizontal lines, while the mean is represented by a thick black horizontal line. The estimated density of the distribution is illustrated by the shaded colour. The dashed line corresponds to the mean for the control conditions. * indicates a significant difference compared to control conditions (Tukey HSD, *p* < 0.05). For each treatment and genotype combination, *n* > 86 in at least 3 independent experiments.

Fig. S4. *sks3* and *sku5* do not suppress the high lateral root density in *max2*

Lateral roots per cm of primary roots in 9-d-old seedlings. Data are shown as mean ± SE. For each genotype, *n* > 51 plants grown in 5 separate experiment. Letters indicate significant differences (Tukey HSD, *p* < 0.05).

Fig. S5. *sks3* does not suppress low germination rate in *max2* Seeds were germinated on 0.8% (w/v) agar plates and germination rate was scored after 72h. Data are shown as mean ± SE, for 10 batches of seeds each batch holding > 80 seeds.• Indicates significant difference compare to the wild type (Tukey HSD, *p* < 0.1).

Fig. S6. Root diameter of *max2-7* and *kai2-2* plants was lower than that of wild type. Letters indicate statistical significance at the 1% level (Tukey, HSD). Data shown as mean ± SE, *n* > 36 per genotype in a total of 5 experiments.

